# Structural characterization of two nanobodies targeting the ligand-binding pocket of human Arc

**DOI:** 10.1101/2023.09.06.556498

**Authors:** José M. Godoy Muñoz, Lasse Neset, Sigurbjörn Markússon, Sarah Weber, Oda C. Krokengen, Aleksi Sutinen, Eleni Christakou, Andrea J. Lopez Moreno, Clive R. Bramham, Petri Kursula

## Abstract

The activity-regulated cytoskeleton-associated protein (Arc) is a complex regulator of synaptic plasticity in glutamatergic neurons. Understanding its molecular function is key to elucidate the neurobiology of memory and learning, stress regulation, and multiple neurological and psychiatric diseases. The recent development of anti-Arc nanobodies has promoted the characterization of the molecular structure and function of Arc. This study aimed to validate two anti-Arc nanobodies, E5 and H11, as selective modulators of the human Arc N-lobe (Arc-NL), a domain that mediates several molecular functions of Arc through its peptide ligand binding site. The structural characteristics of recombinant Arc-NL-nanobody complexes were solved at atomic resolution using X-ray crystallography. The structures revealed that both anti-Arc nanobodies specifically bind to the multi-peptide binding site of Arc-NL. Isothermal titration calorimetry showed that the Arc-NL-nanobody interactions occur at nanomolar affinity, and that the nanobodies can displace a TARPγ2-derived peptide from the Arc-NL binding site. Thus, both anti-Arc-NL nanobodies can be used *in vitro* as competitive inhibitors of endogenous Arc ligands. Differences in the CDR3 loops between the two nanobodies indicate that the spectrum of short linear motifs recognized by the Arc-NL should be expanded. We provide a robust biochemical background to support the use of anti-Arc nanobodies in attempts to target Arc-dependent synaptic plasticity. Function-blocking anti-Arc nanobodies could eventually help unravel the complex neurobiology of synaptic plasticity and help diagnose and treat Arc-associated synaptic disorders.

## Introduction

The mammalian brain is a complex integrator of multisensory information. It can coordinate behavioral responses and promote environmental adaptation. Behavioral assays have highlighted the essential role of memory and learning in survival, as they mediate the integration, stabilization and retrieval of new information, thereby guiding behavior in an adaptative manner [1]. Concurrently, neurobiological studies have shown that memory and learning are mediated by the reorganization of synaptic circuits, where individual neurons selectively modulate the strength of their synapses through synaptic plasticity [2–5].

Plastic changes in synaptic strength involve tightly regulated molecular pathways that can rapidly respond to local variations in neuronal activity [6]. A major regulator of synaptic plasticity is the activity-dependent cytoskeleton-associated protein (Arc). Arc is predominantly expressed in neocortical and hippocampal non-GABAergic CaMKIIα-positive glutamatergic neurons of rats [7]. Hence, Arc has been associated with the regulation of long-term memory and learning in the mammalian brain [8–10]. Arc is essential for the consolidation of different types of memory, playing a key role in the stabilization of memory-associated neural circuitry [11–13]. Furthermore, Arc has been associated with stress regulation [14, 15], sleep homeostasis [16, 17], and Alzheimer’s disease [18–22], as well as other neurological and psychiatric disorders [23–25].

The involvement of Arc in physiological and pathological processes has led to an increased interest in the structure and molecular mechanisms of Arc. Arc is mainly involved in three types of synaptic plasticity: N-methyl-D-aspartate receptor (NMDAR)-dependent long-term potentiation (LTP), metabotropic glutamate receptor (mGluR)-dependent long-term depression (LTD) and synaptic scaling [8, 23]. Therefore, Arc can increase or reduce the synaptic strength of specific synapses (in NMDAR-dependent LTP and mGluR-dependent LTD) and of all the excitatory synapses of a neuron (in synaptic scaling). These molecular pathways are only partially understood.

The molecular functions of Arc are mostly restricted to dendritic spines, where it can regulate the endocytosis of α-amino-3-hydroxy-5-methyl-4-isoxazolepropionic acid receptors (AMPAR) during mGluR-dependent LTD and synaptic scaling [26, 27] and actin polymerization during NMDAR-dependent LTP [28]. The functional complexity of Arc is highlighted by its ability to regulate gene transcription in the nucleus [29, 30] and to form different reversible oligomeric states [31, 32]. Arc can oligomerize into viral-like capsids containing *Arc* mRNA that can be transferred to neighboring cells [33]. This oligomeric state is explained by the retroviral origin of Arc as a repurposed retrotransposon, but the molecular mechanisms underlying capsid formation by mammalian Arc are unknown [33, 34]. Overall, the variety of interrelated molecular functions emphasizes the need to further characterize Arc on a structural and functional level.

Structural biology has provided considerable insights into the functional complexity of Arc, which can be divided into two main protein domains with different functions (Fig 1). A dimerization motif has been traced to the Arc N-terminal domain (NTD) [35], and a peptide binding site was located in the Arc N-lobe (Arc-NL). This domain mediates the interaction between Arc and the transmembrane AMPAR regulatory protein γ2 (TARPγ2, also known as stargazin), promoting the endocytosis of AMPAR [34].

**Fig 1.**
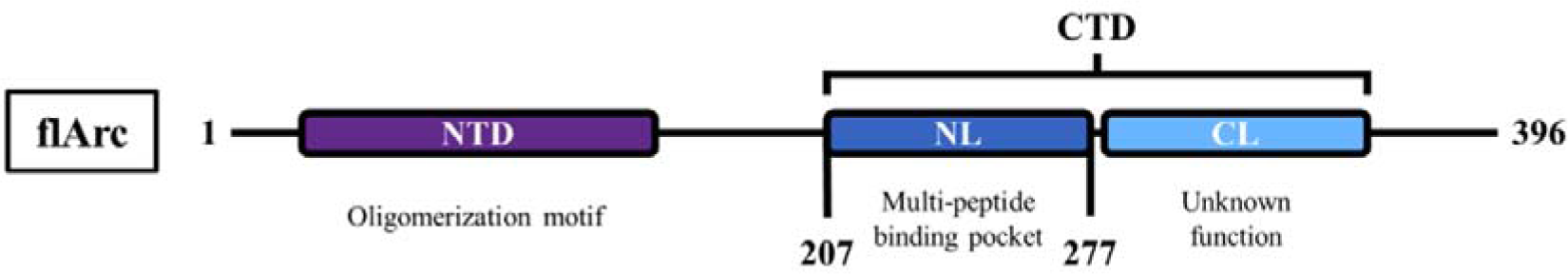
Domains of full-length Arc. Arc is constituted by two main domains: the N-terminal domain (NTD) and the C-terminal domain (CTD). Arc-NTD contains the oligomerization motif, while the Arc-CTD can be further divided into the N-lobe (NL) and the C-lobe (CL). Arc-NL contains a multi-peptide binding pocket that mediates several of the molecular functions of Arc.

The high-resolution structure of full-length Arc (flArc) remains unknown, as the molecular flexibility of Arc hinders its crystallization [36]. To face this problem, anti-Arc nanobodies have been developed and used as crystallization chaperones for the rat and human Arc C-terminal domain (Arc-CTD) [37]. Isothermal titration calorimetry (ITC) revealed that the anti-Arc nanobody H11 can displace a TARPγ2-derived peptide from the multi-peptide binding pocket of flArc, suggesting a role for this nanobody as a functional modulator of Arc function [37]. This finding has prompted an increased interest in anti-Arc nanobodies that selectively bind Arc-NL, as it mediates several essential functions for the regulation of synaptic plasticity, and as the binding event could affect full-length Arc conformation and oligomeric state.

To support the potential applications of anti-Arc-NL nanobodies, we characterized the structural and functional properties of the protein complexes formed between Arc-NL and the anti-Arc nanobodies E5 and H11. Both nanobodies bound to the multi-peptide binding pocket of Arc-NL, which validates them as possible functional modulators of Arc-NL. Nanobody E5 bound flArc simultaneously with nanobody C11, known to target the Arc C-lobe [37, 38]. These findings open a path towards using anti-Arc-NL nanobodies to characterize the different structural states and complex molecular functions of the Arc protein.

## Materials and Methods

### Materials

All chemicals were acquired from Sigma-Aldrich (Merck Group; Massachusetts, US) and all other materials from Thermo Fisher Scientific (Massachusetts, US), unless otherwise specified. For crystal picking, 50-200 µm MicroLoops LD (MiTeGen, Ithaca, New York, US) were used. The modified TARPγ2 peptides were obtained from GenScript Biotech (Piscataway, New Jersey, US).

### Expression constructs

Five recombinant protein constructs were used in this study: anti-Arc nanobody E5, anti-Arc nanobody H11, Arc-NL (Arc_207-277_; identical for human, rat and mouse Arc-NL), rArcFL-7A full-length dimer, and CTdt (Arc-NL+Arc-CL). Anti-Arc nanobodies H11 and E5 were generated in alpacas as described [37, 38] and provided by Nanotag Biotechnologies (Göttingen, Germany) in prokaryotic pNT1433 expression vectors. The recombinant nanobodies carried an N-terminal histidine-tag followed by a tobacco etch virus (TEV) protease cleavage site (His_6_-TEV-E5/H11). The Arc-NL construct was obtained as a prokaryotic pETMBP_1a expression vector fused with maltose binding protein (His _6_-MBP-TEV-NL) [36]. The full-length rat Arc construct containing a poly-Ala mutation in residues 113-119 (rArcFL-7A) was expressed with an N-terminal fusion (His _6_-MBP) in a pHMGWA vector [35]. The human Arc CTD without its C-terminal tail (Arc-CTdt, residues 206-361) was expressed using the pTH27 vector [36].

### Recombinant protein expression and purification

All recombinant proteins were expressed in *E. coli* and purified as described [37, 39]. Briefly, all proteins were expressed in BL21(DE3) *E. coli* and purified using nickel-nitrilotriacetic acid (Ni-NTA) affinity chromatography. After proteolytic cleavage of the tag and dialysis, a negative Ni-NTA affinity chromatography was conducted to remove His-tagged contaminants. For rArcFL-7A, an amylose affinity chromatography step was additionally included. For size exclusion chromatography (SEC), the sample was concentrated and applied to a HiLoad 16/600 Superdex 75 pg or a HiLoad 16/60 Superdex 200 pg column (GE Healthcare, Illinois, US). The purity of each protein fraction was assessed through SDS-PAGE; the purest fractions were pooled, concentrated, and snap-frozen for storage at −80 °C. Protein concentration was measured with a NanoDrop 2000 spectrophotometer (Thermo Scientific, Massachusetts, US) or with an Abbemat Performance 500 Refractometer (Anton Paar GmbH, Graz, Austria).

### Dynamic Light Scattering (DLS)

DLS was performed using Zetasizer Nano ZS (Malvern Panalytical, Malvern, UK). Samples were filtered using an Ultrafree-MC-GV Centrifugal Filter (Durapore-PVDF, 0.22 μm) (Merck KGaA, Darmstadt, Germany), diluted to 2 mg/ml of protein in 40 μl and loaded into a quartz precision cell (light path: 3×3 mm; Z-height: 8.5 mm) (Hellma, Müllheim, Germany). DLS measurements were taken at 4 °C with a pre-incubation period of 60-120 s. Three measurements of each sample were taken (12 runs/measurement, 10 s/run).

### Circular dichroism (CD) spectroscopy

CD spectroscopy was performed using a Jasco J-810 spectropolarimeter (Jasco, Tokyo, Japan). Before each experiment, the samples were dialyzed overnight against 10 mM phosphate buffer (pH 7.5) and filtered with Ultrafree-MC-GV Centrifugal Filters (Durapore-PVDF, 0.22 μm) (Merck KGaA, Darmstadt, Germany). Filtered samples containing 0.15-0.20 mg/ml of protein were loaded into a 1-mm quartz cuvette (Hellma, Müllheim, Germany). CD spectra were measured at 20 °C in continuous scanning mode using the following parameters: spectral width = 185-280 nm; standard sensitivity = 100 mdeg; scan speed = 50 nm/min; data pitch = 1 nm; 4 s response; bandwidth = 1 nm. Each run consisted of the accumulation of four individual spectra, and buffer spectra were subtracted.

The CD spectra were analyzed using CDToolX (version 2.01, Birkbeck College, University of London) [40]. The raw units (mdeg) were transformed into Δε (M^-1^ cm^-1^).

### Synchrotron radiation circular dichroism (SRCD) spectroscopy

High-resolution SRCD spectra were measured on the AU-CD beamline at the synchrotron storage ring ASTRID2 (ISA, Aarhus, Denmark), using 0.15-0.20 mg/ml protein samples dialyzed and filtered as above. 30 μl of sample were loaded into a 0.1-mm pathlength closed circular cuvette (Hellma, Müllheim, Germany). SRCD spectra were measured at 25 °C using a wavelength range of 170-280 nm. Each run consisted of the accumulation of six individual spectra.

To determine the denaturation midpoint (T _m_) of each protein sample, temperature scans were carried out by measuring the CD spectrum of each sample between 24.2-84.5 °C (heating rate 2.5 °C/min). The ellipticity values at 208 nm (for Arc-NL, Arc-NL-E5 and Arc-NL-H11) and 190 nm (for E5 and H11) were chosen as indicators of folding state. Assuming a two-state denaturation process (Folded protein [F] → Unfolded protein [U]), the ellipticity values were transformed to represent the fraction of unfolded protein [41]. Then, the T _m_ of each protein was calculated by fitting the data to a sigmoidal curve using the Boltzmann sigmoidal function in GraphPad Prism (v.9.5.1).

### CD spectral deconvolution

Individual CD and SRCD spectra were averaged in CDToolX [40]. The averaged datasets were uploaded to BeStSel [42]. Deconvolution was performed using a scale factor of 1 for several wavelength ranges. The deconvolution profile with the lowest associated root-mean-squared deviation factor (RMSD) was chosen as the optimal profile [43, 44].

### Protein crystallization

Protein crystallization was carried out using the sitting drop method for anti-Arc nanobody H11, Arc-NL in complex with E5, and Arc-NL in complex with H11. Multiple crystallization conditions were tested using the commercial screens JCSG+ and PACT Premier (Molecular Dimensions, Sheffield, UK) at two temperatures (8 °C and 20 °C) in 96-well sitting drop iQ plates (STP Labtech, Melbourn, UK). The Arc-NL-nanobody complexes were formed by directly mixing equimolar amounts of each protein (for Arc-NL-E5) or a 1.5-fold excess of nanobody (for Arc-NL-H11). Protein samples (6-9 mg/ml) were plated using a Mosquito LCP crystallization robot (SPT Labtech, Melbourn, UK); for each condition, three drops (300 nl) with different protein:mother liquor ratios (2:1, 1:1 and 1:2) were deposited next to a 70 μl reservoir containing the precipitant well solution.

H11 (8.7 mg/ml, TBS buffer) crystallized at 8 °C in a 300-nl drop with a 1:1 protein:well solution ratio after ∼30-60 days. The well solution contained 0.1 M Bis-Tris (pH 5.5) and 25% w/v PEG 3350. The crystal was not cryoprotected during cryocooling.

The Arc-NL-H11 complex (4.2 mg/ml H11, 1.8 mg/ml Arc-NL, TBS buffer) crystallized at 20 °C in a 300-nl drop with a 1:2 protein:well solution ratio after ∼14 days. The well solution contained 0.1 M potassium phosphate/citrate pH 4.2, 0.2 M NaCl and 20% w/v PEG 8000. The crystal was not cryoprotected.

The Arc-NL-E5 complex (4.1 mg/ml E5, 2.8 mg/ml Arc-NL, TBS buffer) crystallized at 20 °C in a 300-nl drop with a 1:1 protein:well solution ratio after ∼30 days. The well solution contained 0.1 M Tris pH 8, 0.01 M ZnCl _2_ and 20% w/v PEG 6000. The crystal was cryoprotected using well solution supplemented with 25% glycerol.

### X-ray diffraction data collection and structure determination

Data collection took place on the P11 beamline at the German Electron Synchrotron (DESY, Hamburg, Germany) [45]. The data were collected at 100 K on an Eiger 2X 16M detector with a 50×50 μm^2^ focused beam at 25-35% transmission. Images were taken using an oscillation range of 0.1° and 10 ms exposure per frame. The data were processed with XDS [46] and evaluated using Xtriage [47].

The crystallographic phases were solved via MR using the structures of anti-Arc nanobody E5 (PDB ID: 7R20) [37] and the Arc-NL bound to a stargazin peptide (PDB ID: 6TNO) [39]. Refinement was carried out in phenix.refine [48] and manual model building in Coot [49]. Data processing and refinement statistics are in Table 1. Structure validation was conducted using MolProbity [50]. The analysis of protein interfaces was carried out using PISA [51]. Molecular representations and structural analyses of protein interfaces were done using UCSF Chimera [52] and PyMOL.

**Table 1.** X-ray diffraction data and refinement statistics. Data in parentheses correspond to the highest-resolution shell.

|  |  |  |  |
| --- | --- | --- | --- |
| Protein | Arc-NL-H11 | Arc-NL-E5 | H11 |
| Beamline | P11/DESY | P11/DESY | P11/DESY |
| Wavelength (Å) | 0.83 | 1.03 | 1.03 |
| Resolution range (Å) | 50–1.02 (1.05–1.02) | 50–1.50 (1.54–1.50) | 50–1.25 (1.28–1.25) |
| Space group | P2<sub>1</sub> | P22<sub>1</sub>2<sub>1</sub> | P4<sub>3</sub>2<sub>1</sub>2 |
| Unit cell parameters | a = 41.82 Å<br>b = 52.07 Å<br>c = 44.22 Å<br>α = γ = 90°<br>β = 109.89° | a = 34.50 Å<br>b = 99.48 Å<br>c = 138.32 Å<br>α = β = γ = 90° | a = b = 57.17 Å<br>c = 145.64 Å<br>α = β = γ = 90° |
| Completeness (%) | 95.2 (79.9) | 99.5 (99.9) | 99.9 (99.7) |
| Mean I/σ(I) | 13.0 (0.8) | 15.0 (0.7) | 13.1 (0.6) |
| Wilson B-factor (Å<sup>2</sup>) | 15.4 | 37.9 | 25.8 |
| R<sub>cryst</sub> (%) | 5.4 (201.7) | 4.9 (287.9) | 7.1 (367.9) |
| CC<sub>1/2</sub> (%) | 99.9 (31.6) | 99.9 (31.7) | 99.9 (28.2) |
| R<sub>work</sub> (%) | 12.9 | 14.5 | 18.5 |
| R<sub>free</sub> (%) | 16.0 | 19.3 | 23.2 |
| RMSD bond lengths(Å) | 0.020 | 0.011 | 0.018 |
| RMSD bond angles (°) | 1.58 | 1.10 | 1.70 |
| Ramachandran favored (%) | 98.4 | 97.3 | 99.60 |
| Ramachandran outliers (%) | 0 | 0 | 0 |
| MolProbity score/percentile | 1.25/89<sup>th</sup> | 1.19/97<sup>th</sup> | 1.22/95<sup>th</sup> |
| PDB code | 8QF4 | 8QF5 | 8QF3 |

### Size exclusion chromatography – multi-angle light scattering (SEC-MALS)

SEC-MALS was performed to estimate the oligomeric state and stoichiometry of the rArcFL7A dimer and its nanobody complexes. SEC was performed on a HiLoad 16/60 Superdex 200 pg SEC column (GE Healthcare) equilibrated with 20 mM Tris pH 7.5, 150 mM NaCl and 0.5 mM TCEP using a Shimadzu Prominence-i LC-2030C 3D HPLC unit (GMI, MN, USA) with an LC-2030/2040 PDA UV detector (Shimadzu, Kyoto, Japan). The system was calibrated using bovine serum albumin shortly before running the samples. 50 μg of each protein complex in the running buffer were injected at a flow rate of 0.5 ml/min and light scattering and refractive index were recorded using a miniDAWN TREOS detector (Wyatt Technologies, CA, USA) and a RefractoMax 520 refractometer (Wyatt Technologies). Data collection and SEC-MALS analysis were carried out using ASTRA 6.1 (Wyatt Technologies).

### Analytical SEC

To see if the Arc CTdt (C-terminal domain lacking the disordered tail) construct was in fact able to bind two nanobodies simultaneously and to make sure the complex peak would not overlap with the excess nanobody peaks, a small-scale analytical SEC was run. For each run, 3 nmol CTdt were used, and the nanobodies H11 and C11 were added in 1.3-fold excess (3.9 nmol). SEC was run on a Superdex 75 increase 10/300 GL column (GE Healthcare), at a flow rate of 0.6 ml/min. The running buffer was 20 mM Tris pH 7.4, 150 mM NaCl.

### Small-angle X-ray scattering

In SEC-SAXS, the sample is eluted from the gel filtration column directly into the X-ray beam. Arc and the nanobodies were combined in a 1:1.3 protein-to-nanobody ratio to ensure all the Arc protein is bound to the nanobodies. The proteins were gently mixed and incubated on ice for 1 h. After incubation, the proteins were filtered using an Ultrafree 0.22 μm centrifugal filter (Merck Millipore) to remove any aggregation. For SEC-SAXS, Agilent Bio SEC-3 Column (Agilent Technologies, CA, USA) or HiLoad 16/60 Superdex 200 pg SEC column (GE Healthcare) were used, with 20 mM Tris pH 7.5, 150 mM NaCl and 0.5 mM TCEP as running buffer.

SEC-SAXS data collection was performed twice. First on the CoSAXS beamline [53] at MAX IV (Lund, Sweden) and again on the SWING beamline [54] at the SOLEIL synchrotron (Gif-sur-Yvette, France). The data shown here are from SOLEIL, due to better signal-to-noise ratio and lack of radiation damage. Buffer subtraction and frame selection were performed in CHROMIXS [55], primary analysis in PRIMUS [56] and distance distribution function analysis using GNOM [57]. *Ab initio* model building was done with DAMMIN [58] and GASBOR [59].

### Functional assay using ITC

ITC was performed with a MicroCal iTC200 instrument (Malvern Panalytical, Malvern, UK). The temperature was set to 20 °C, using a reference power of 5 μcal/s, a stirring speed of 1000 rpm, and a filter period of 5 s. For titration of the nanobodies into Arc-NL, 15 μM E5/H11 (titrant) and 1.5 µM Arc-NL (titrate), both diluted in TBS (20 mM Tris-HCl pH 7.5, 150 mM NaCl), were used. Each titration consisted of a 0.5 μL initial injection followed by 18 injections (2 μL, 4 s each; separated by 120 s). Four high-quality replicates were obtained for each nanobody.

Titration of FITC-labelled TARPγ2 peptide (P1) into Arc-NL (positive control) was conducted with 2 mM P1 (titrant) and 200 µM Arc-NL (titrate), both diluted in TBS. Three sequential titrations were carried out to ensure that saturation was reached at the end of the assay; each titration consisted of a 0.5 μL initial injection followed respectively by 18, 10, and 5 injections (2 μL, 4 s each, separated by 120 s). The three individual data files were concatenated using the ConCat32 tool (provided by the manufacturer) into a single datafile.

Titration of FITC-labelled reverse TARPγ2 peptide (P2) into Arc-NL (negative control) was conducted with 0.5 mM P2 (titrant) and 50 µM Arc-NL (titrate), both diluted in TBS. The titration consisted of one 0.5 μL initial injection followed by 9 injections (4 µl, 8 s each; separated by 240 s).

For the displacement assays, 100 μM E5/H11 (titrant), and a titrate solution of 10 µM Arc-NL saturated with 100 μM P1, were used. Both solutions were diluted in TBS. Each titration consisted of a 0.5 μL initial injection followed by 18 injections (2 μL, 4 s each; separated by 120 s).

### ITC data analysis

Data analysis was carried out following a pre-established protocol based on three inter-connected programs [60]. Automatic peak integration and control subtraction were conducted in NITPIC (v. 2.0.7) [61]. The integrated data were fitted to a sigmoidal curve using a 1:1 binding model in SEDPHAT (v. 14.1) [62]. SEDPHAT was also used to calculate and subtract (1) the heats of dilution and (2) the heats of ionization of the buffer. After fitting, best-fit values and their 68.3% confidence interval (± σ) were obtained for the binding enthalpy (ΔH) and association constant (K_A_ = 1/K_D_). From these values, the best-fit values for binding entropy (ΔS) and free energy (ΔG) were calculated by the software. The graphs shown in S1 Fig were plotted using GUSSI (v. 1.7.2) [63].

### Sequence homology search of the CDR3s of anti-Arc-NL nanobodies

Sequence homology searches were carried out in Protein BLAST [64] using the CDR3 of both anti-Arc-NL nanobodies E5 and H11 as query sequences. The results were restricted to three target species: *Homo sapiens* (taxid:9606), *Rattus norvegicus* (taxid:10116) and *Mus musculus* (taxid:10090). The non-redundant protein sequence database was used. Due to the short length of the query sequence, only matches with an E-value < 100 were studied, and only those with an E-value ≤ 10 were considered relevant. After extracting the sequences, sequence alignment between each CDR3 and selected sequences was performed using Clustal Omega [65] and visualized in Jalview [66].

## Results and Discussion

### Protein characterization

For interaction assays and high-resolution structural studies, Arc-NL and the anti-Arc nanobodies E5 and H11 were purified in high yield. DLS shows that the three samples are monodisperse (Fig 2A). None of the purified proteins show a tendency for aggregation, and their hydrodynamic radii (R_H_) correspond to their respective molecular weights (i.e., R _H_-Arc-NL [8.9 kDa] < R _H_-E5 [12.8 kDa] < R _H_-H11 [14.1 kDa]). In addition, rArcFL-7A and Arc-CTdt were purified as previously shown [37].

**Fig 2.**
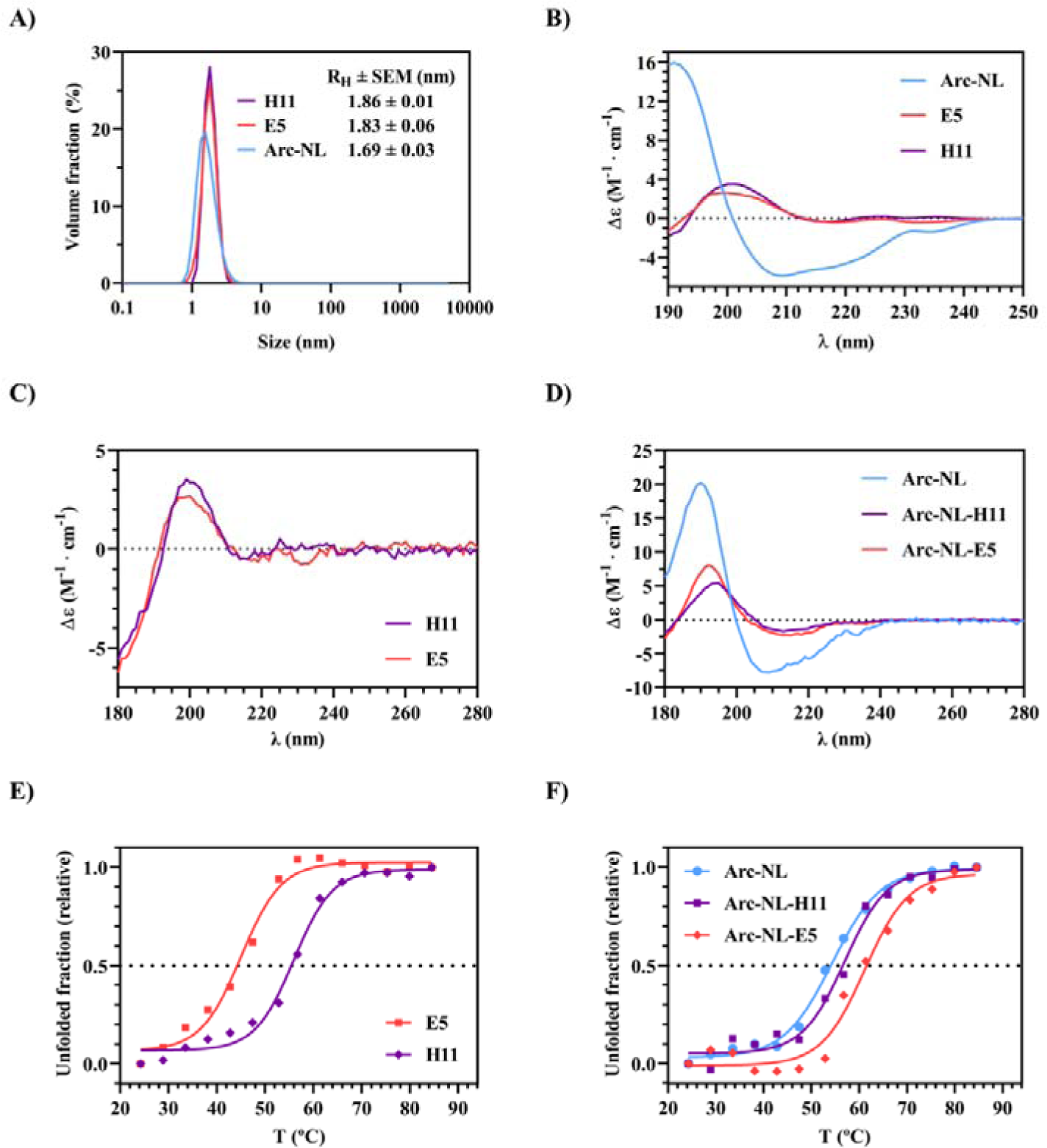
Characterization of Arc-NL, E5 and H11. (A) DLS analysis of Arc-NL, E5 and H11. (B) CD spectroscopy of Arc-NL, E5 and H11. (C) SRCD spectroscopy of H11 and E5. (D) SRCD spectroscopy of Arc-NL, and the Arc-nanobody complexes Arc-NL-H11 and Arc-NL-E5. (E) Thermal denaturation curves based on SRCD of H11 and E5. (F) Thermal denaturation curves based on SRCD of Arc-NL and the Arc-nanobody complexes Arc-NL-H11 and Arc-NL-E5.

CD spectra for Arc-NL, E5 and H11 show that all proteins are likely to be correctly folded (Fig 2B-2D). For Arc-NL, the spectra show that the structure is mainly α-helical (Fig 2B and 2D) [67]. For both nanobodies, the ellipticity signal is weak above 210 nm, indicating structural features canceling each other in the spectrum (Fig 2B and 2C). The positive peak at 200 nm is a strong indicator that both nanobodies consist mostly of β-sheets (Fig 2B and 2C). The presence of these secondary structure components and the lack of negative ellipticity at 195 nm (characteristic of disordered proteins) [67] indicate that all proteins are correctly folded. The SRCD spectra of both Arc-nanobody complexes show a negative peak around 208 nm, further indicating that Arc-NL is folded in solution with both nanobodies. The deconvolution profiles of all CD and SRCD spectra are shown in Table 2.

**Table 2.**
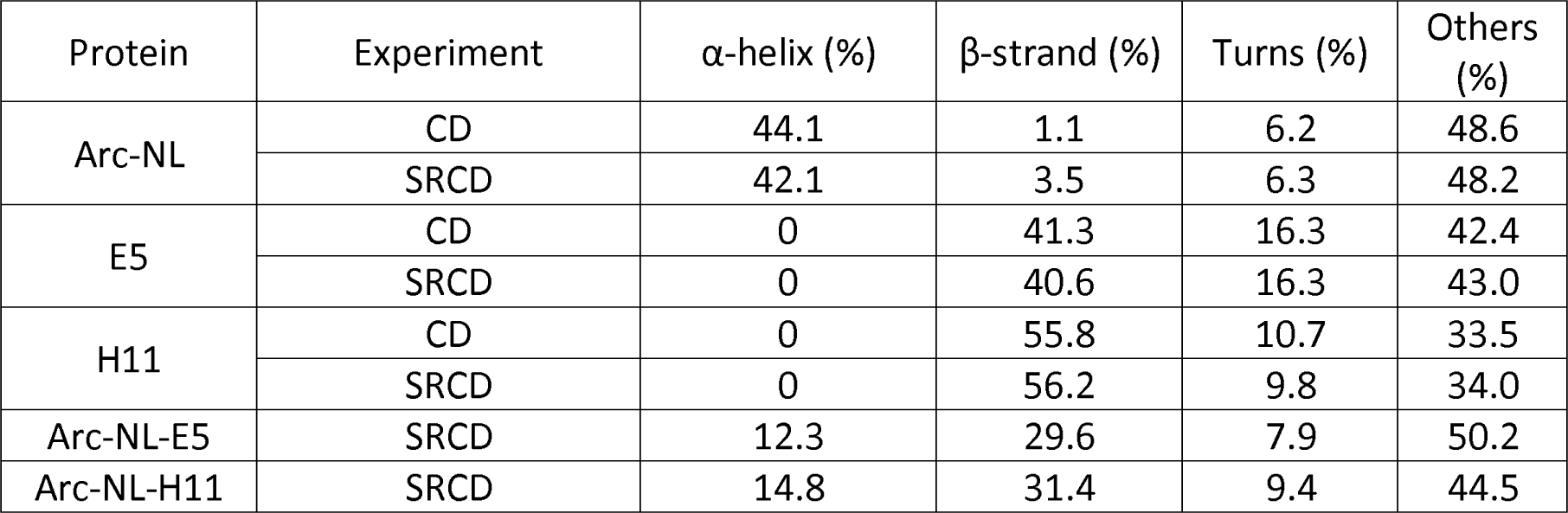
Deconvolution profiles of CD/SRCD spectra. Deconvolution was done using BeStSel of Arc-NL, E5, H11, and Arc-nanobody complexes Arc-NL-E5 and Arc-NL-H11. The BeStSel algorithm divides the deconvoluted signals into four main categories of protein secondary structures: α-helices, β-strands, turns and others.

BeStSel was chosen as the optimal deconvolution software to analyze the nanobody CD spectra, as it is sensitive to different conformations of β-strands [42]. The deconvolution supports a predominance of α-helices in Arc-NL and β-strands in the nanobodies [34, 37]. Altogether, DLS and CD/SRCD spectroscopy show that the recombinant proteins are monodisperse and correctly folded.

SRCD spectroscopy was further used to study the thermal stability of the proteins and complexes. Both complexes have a higher T _m_ than any of the proteins alone (Fig 2E and 2F; Table 3). The Arc-NL-E5 complex shows the highest increase in T_m_ of 16.1 °C. The data show an increase in thermal stability when the nanobodies bind to Arc, and the presence of E5 causes stronger stabilization. The results hint towards at least partially different binding modes for the two nanobodies.

**Table 3.** Thermal denaturation midpoints (T_m_) of Arc-NL, E5, H11, and Arc-nanobody complexes Arc-NL-E5 and Arc-NL-H11. The best-fit V50 value of the Boltzmann function applied to the thermal denaturation curve is shown together with its 95 % confidence interval.

| Protein | $T_m$ (°C) | 95 % CI (°C) |
| --- | --- | --- |
| Arc-NL | 54.33 | (53.29, 55.39) |
| E5 | 45.07 | (42.72, 47.00) |
| H11 | 55.97 | (54.45, 57.36) |
| Arc-NL-E5 | 61.16 | (58.82, 63.93) |
| Arc-NL-H11 | 56.97 | (55.10, 58.82) |

### Both E5 and H11 interact with the peptide binding pocket of Arc-NL

Crystallization was successful for Arc-NL-H11, Arc-NL-E5 and unbound H11, and structures could be refined at atomic resolution (1.02 Å for Arc-NL-H11, 1.5 Å for Arc-NL-E5 and 1.33 Å for H11). In addition, the crystal structure of apo-E5 had been solved before [37]. The crystals that underwent X-ray diffraction are shown in Fig 3A, together with the refined structures (Fig 3B). See Table 1 for detailed X-ray diffraction data and refinement statistics.

**Fig 3.**
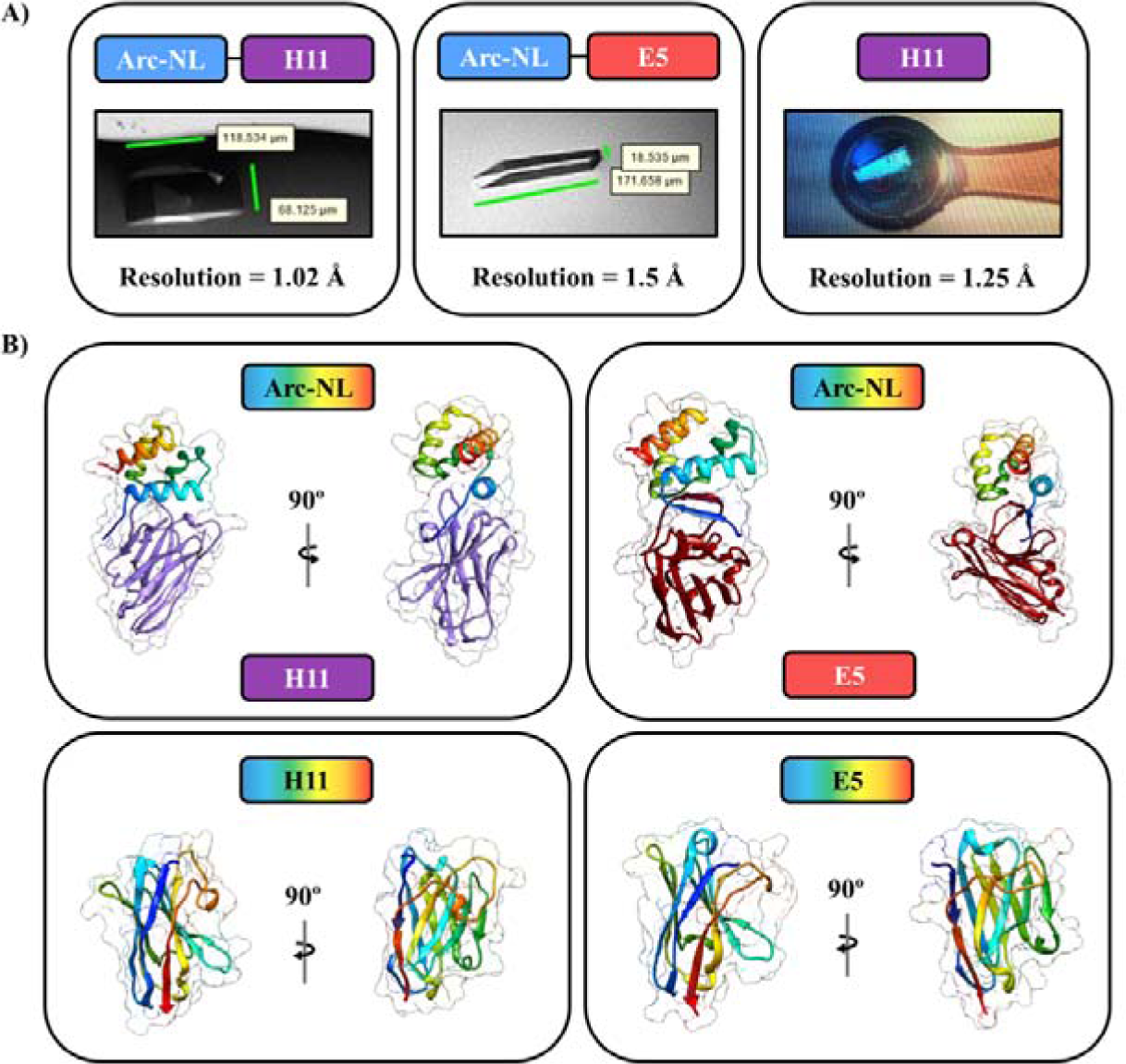
Protein crystals and refined structures for Arc-NL-H11, Arc-NL-E5 and H11. (A) Crystals of Arc-NL-H11, Arc-NL-E5 and unbound H11. (B) The structures of Arc-NL-H11 (top left), Arc-NL-E5 (top right) and unbound H11 (bottom left). The apo E5 structure (bottom right) was previously solved (PDB ID: 7R20) [37] and is shown for comparison with the Arc-NL-E5 complex.

A structural analysis of the Arc-nanobody complexes reveals that E5 and H11 both bind the multi-peptide binding site of Arc-NL (Fig 4A). When binding Arc-NL, both E5 and H11 undergo conformational changes in their CDRs (Fig 4B). It appears that the CDR3 of E5 forms a β-sheet with the Arc-NL N-terminal loop when the complex is formed (Fig 4A and 4B), similarly to the TARPγ2 ligand peptide [34, 39]. Hence, the conformation of the N terminus of Arc-NL is different in the E5 and H11 complexes.

**Fig 4.**
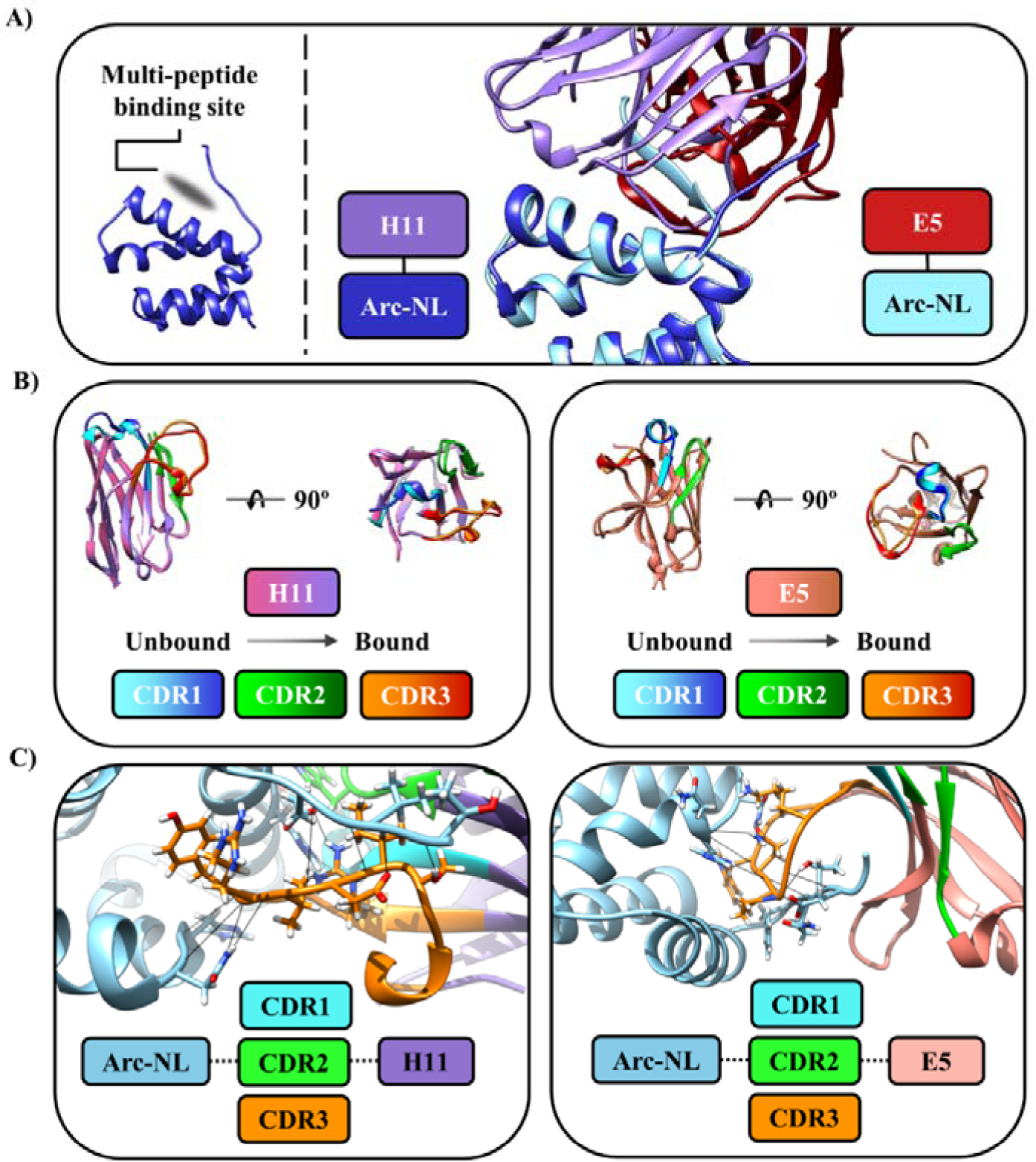
Structural analysis of Arc-nanobody complexes. (A) Both E5 and H11 bind to the multi-peptide binding pocket of Arc-NL. (B) Overlay of bound and unbound H11 and E5, showing the conformational changes that each nanobody displays when binding Arc-NL. (C) Intermolecular interactions between each nanobody and Arc-NL. The CDR3 of both nanobodies is the main contributor to the protein complex formation.

Table 4 summarizes the interfaces of the complexes. For both Arc-nanobody complexes, there is one main interface contributing to the complex formation that is estimated to lead to a solvation free energy gain (ΔG _s_ < 0) in an interaction-specific manner (i.e., not as a consequence of crystal packing), represented by P < 0.5 [51]. For both nanobodies, CDR3 is the main contributor to the complex formation (Fig 4C). The interface between H11 and Arc-NL involves CDR1, 2, and 3 of H11 (Fig 4C), while binding between E5 and Arc-NL only involves the CDR3 of the nanobody (Fig 4D). This is intriguing, as the heat stability increase in the complex was higher for E5; this can be related to the higher number of hydrogen bonds present at the interface for E5 (Table 4).

**Table 4.**
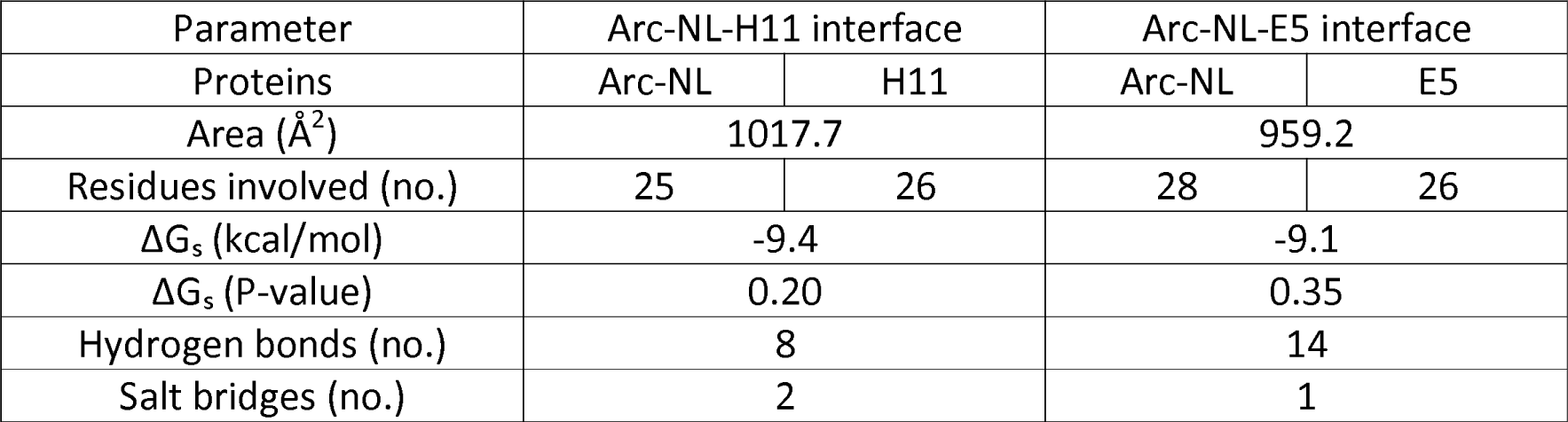
Protein interface analysis of the Arc-NL-H11 and Arc-NL-E5 complexes. Main parameters derived from PDBePISA. The main interface that contributes to the complex formation and its associated parameters are shown.

| Parameter | Arc-NL-H11 interface |  | Arc-NL-E5 interface |  |
| --- | --- | --- | --- | --- |
| Proteins | Arc-NL | H11 | Arc-NL | E5 |
| Area ($\text{\AA}^2$) | 1017.7 | | 959.2 | |
| Residues involved (no.) | 25 | 26 | 28 | 26 |
| $\Delta G_s$ (kcal/mol) | -9.4 | | -9.1 | |
| $\Delta G_s$ (P-value) | 0.20 | | 0.35 | |
| Hydrogen bonds (no.) | 8 |  | 14 |  |
| Salt bridges (no.) | 2 |  | 1 |  |

According to the protein interface statistics, the formation of the Arc-NL-H11 interface appears to be slightly more favorable energetically than for Arc-NL-E5 (ΔG_s_ [Arc-NL-H11] < ΔG_s_ [Arc-NL-E5]). However, the measured T_m_ implies that Arc-NL-E5 is more stable than Arc-NL-H11 (T_m_ [Arc-NL-E5] > _m_T [Arc-NL-H11]). Hence, the two nanobodies bind to overlapping epitopes on Arc-NL with unique modes of interaction, however producing similar affinity.

In addition, the structural study of unbound E5 and H11 provides a potential explanation for the weak ellipticity found above 210 nm in CD/SRCD spectra. E5 contains Trp^38^ that is in close proximity of a disulfide bond formed between Cys^24^ and Cys^97^ [37]. The structure of unbound H11 also shows this feature; Trp^38^ is located next to a disulfide bond (Cys^24^-Cys^98^). Aromatic residues can distort the CD spectra of β-strand-containing proteins by inducing positive ellipticity around 215-225 nm [68, 69], especially if they are located close to disulfide bonds [70]. Therefore, this common nanobody structural feature can explain the observed characteristic CD spectra.

### Nanobodies E5 and H11 displace a TARPγ2-derived peptide from the Arc-NL peptide binding pocket

The thermodynamic properties of the Arc-NL-nanobody complex formation were determined using ITC. The thermodynamic parameters associated with all ITC assays are presented in Table 5. The individual replicates carried out for Arc-nanobody interactions are shown in S1 Fig. Both nanobodies show a similar profile, binding to Arc-NL with high affinity (K_D_ ≈ 1 nM) in an enthalpy-driven process (ΔH ≈ −23 kcal/mol) (Fig 5A), with unfavorable entropy indicative of e.g. ordering of the long CDR3 loop upon binding. Arc-NL shows exothermic binding (K _D_ = 5.3 μM) to a TARPγ2-derived peptide (P1), while binding is not observed when titrating the reverse control peptide (P2) (Fig 5B). A displacement assay was carried out for both nanobodies; each assay consisted of the titration of the nanobody into a solution of Arc-NL saturated with P1 (Fig 5C, Fig 5D, Table 5). Both nanobodies show an apparent reduction of their binding affinity and binding enthalpy. The reduction in binding enthalpy corresponds in both cases to 11-13 kcal/mol, which is similar to the dissociation enthalpy of P1 and Arc-NL (-ΔH = 14.9 kcal/mol). This implies that the dissociation of P1 and Arc-NL is coupled to the association between E5/H11 and Arc-NL, suggesting that both nanobodies can displace the TARPγ2-derived peptide from Arc-NL. In addition to the crystal structures, this is supported by the fact that P1 and the CDR3s of both nanobodies contain the consensus motif for binding to Arc-NL: PxY/W/F [36] (Fig 5E).

**Fig 5.**
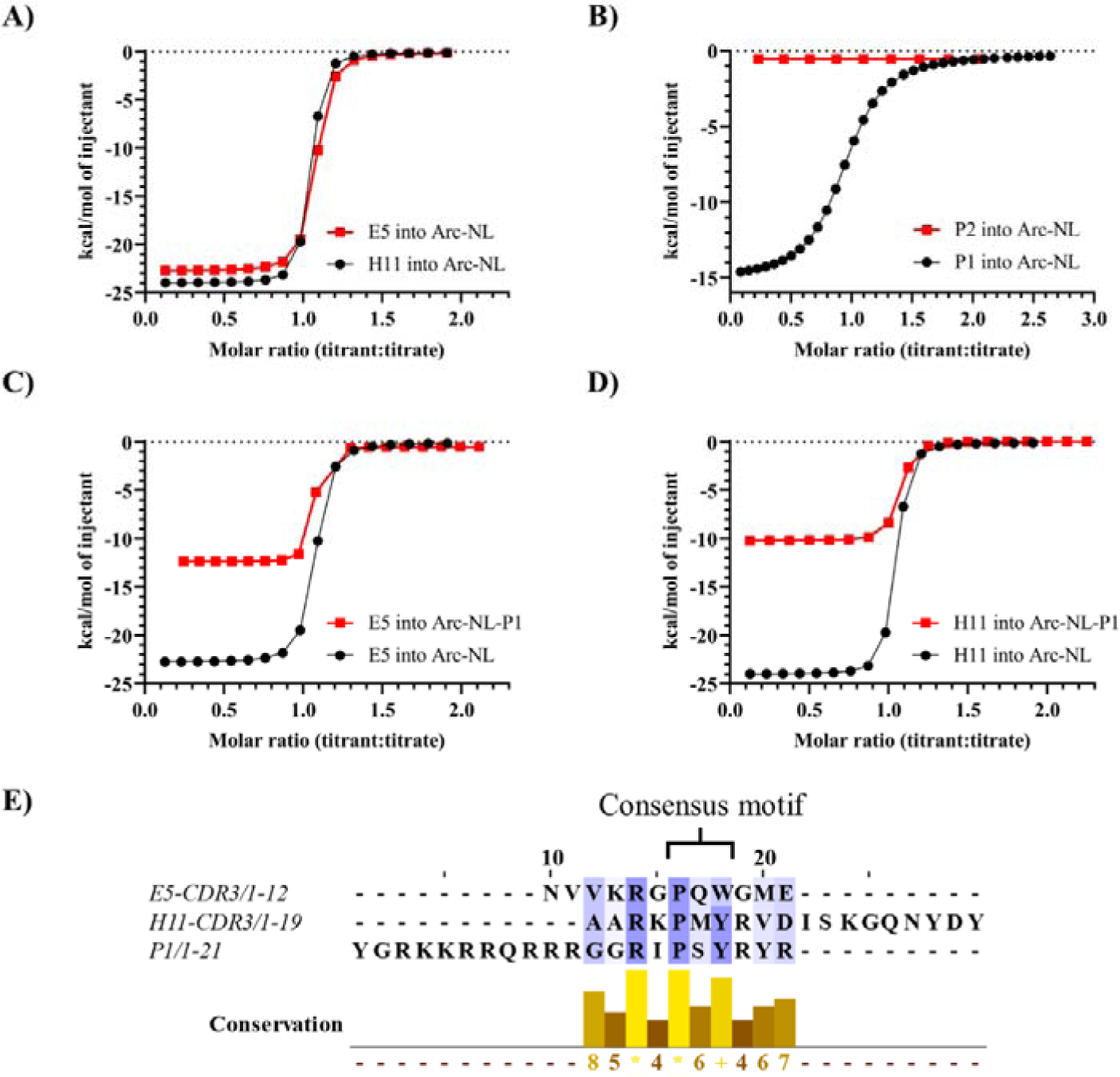
Thermodynamic characterization of the binding dynamics between Arc-NL, TARPγ2-derived peptides, and anti-Arc-NL nanobodies. (A) Titration of E5/H11 into Arc-NL; both nanobodies display a similar thermodynamic binding profile. Only one representative replicate of each assay is shown. (B) Titration of P1/P2 into Arc-NL; binding is selective for P1, while P2 is valid as a negative control. (C) Displacement assay for E5. An apparent reduction in K _D_ and ΔH can be observed. (D) Displacement assay for H11. The apparent reduction of the K _D_ and ΔH values is seen. (E) Multiple sequence alignment of P1 and the CDR3s of H11 and E.

**Table 5.** Thermodynamic parameters. For Arc-nanobody assays, four replicates were analyzed (n = 4); the mean best-fit values ± SEM of their thermodynamic binding properties are displayed. For the P1/P2-Arc-NL assays and the displacement assays, single high-quality replicates were taken. For each individual replicate, the binding affinity and enthalpy are shown alongside their 68.3 % CI (± σ), as they are measures directly calculated from the experimental thermograms by SEDPHAT [62]. Abbreviations: ΔG, free energy of binding; ΔG_App_, apparent free energy of binding; ΔH, binding enthalpy; ΔH _App_, apparent binding enthalpy; CI, confidence interval; K_D_, binding affinity (dissociation constant); K _D-App_, apparent binding affinity (apparent dissociation constant); P1, FITC-labelled TARPγ2-derived peptide; P2, FITC-labelled TARPγ2 reverse peptide.

| Binding assay | $K_D$ (nM) | | $\Delta H$ (kcal/mol) | | $-T\Delta S$ (kcal/mol) | $\Delta G$ (kcal/mol) |
| --- | --- | --- | --- | --- | --- | --- |
| E5 into Arc-NL ($n = 4$) | $0.99 \pm 0.27$ | | $-23.24 \pm 0.40$ | | $+10.97 \pm 0.30$ | $-12.27 \pm 0.22$ |
| H11 into Arc-NL ($n = 4$) | $1.14 \pm 0.56$ | | $-23.48 \pm 0.71$ | | $+11.26 \pm 0.43$ | $-12.22 \pm 0.41$ |
| Binding assay | $K_D$ ($\mu M$) | $K_D - 68.3\% \text{ CI}$ ($\mu M$) | $\Delta H$ (kcal/mol) | $\Delta H - 68.3\% \text{ CI}$ (kcal/mol) | $-T\Delta S$ (kcal/mol) | $\Delta G$ (kcal/mol) |
| P1 into Arc-NL ($n = 1$) | 5.31 | (5.02, 5.62) | -14.86 | (-14.96, -14.77) | +7.79 | -7.08 |
| P2 into Arc-NL ($n = 1$) | $6.92 \cdot 10^{21}$ | - | 0.049 | - | +25.22 | +25.27 |
| Displacement assay | $K_{D-App}$ (nM) | $K_{D-App} - 68.3\% \text{ CI}$ (nM) | $\Delta H_{App}$ (kcal/mol) | $\Delta H_{App} - 68.3\% \text{ CI}$ (kcal/mol) | $-T\Delta S_{App}$ (kcal/mol) | $\Delta G_{App}$ (kcal/mol) |
| E5 into Arc-NL-P1 ($n = 1$) | 6.93 | (1.00, 14.69) | -11.98 | (-12.20, -11.76) | +1.03 | -10.95 |
| H11 into Arc-NL-P1 ($n = 1$) | 10.38 | (2.48, 25.73) | -10.30 | (-10.83, -9.81) | -0.411 | -10.71 |

The above experiments show that binding between TARPγ2 and Arc-NL can be inhibited by anti-Arc nanobodies E5 and H11. Although displacement assays were not performed for other Arc ligands, binding to the Arc-NL peptide binding pocket should be similarly inhibited by E5 and H11 for other endogenous Arc-NL ligands. TARPγ2 stands out as the endogenous Arc-NL ligand with the highest known affinity [39]; therefore, if binding to TARPγ2 is blocked by H11 and E5, these two nanobodies will also displace lower-affinity ligands. Interestingly, the conformation of the Arc-NL N terminus bound to peptides derived from Arc endogenous ligands is strikingly similar to the conformation it adopts when binding E5 (Fig 6). It is possible that conformational differences exist in flArc, when bound to either E5 or H11, as the N terminus of Arc-NL corresponds to the end of the flexible linker region. This may raise possibilities for conformation-specific binding or conformational modulation in the context of flArc, despite overlapping binding sites for the two nanobodies.

**Fig 6.**
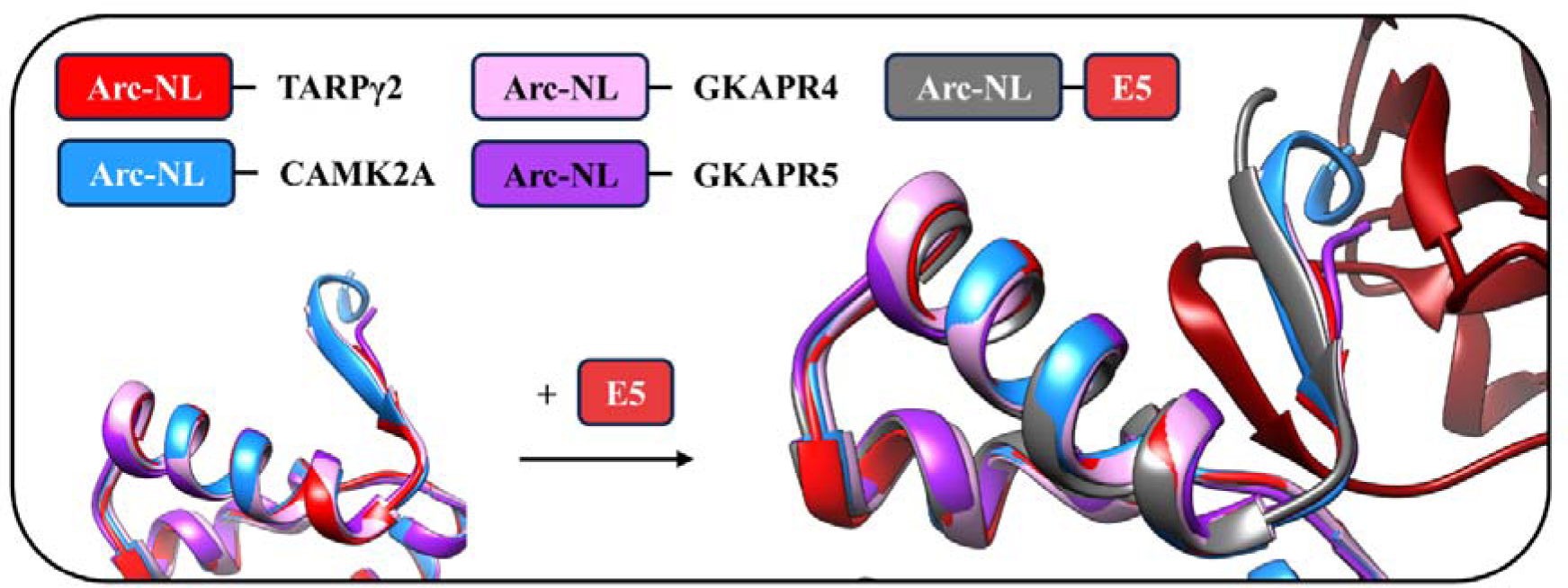
Comparative structural analysis of Arc-NL bound to different peptides derived from endogenous Arc ligands. The Arc N-lobe adopts a similar conformation while interacting with different ligand peptides: TARPγ2 (PDB ID: 6TNO) [39], CAMK2A (PDB ID: 4X3I) [34], GKAPR4 (PDB ID: 6TNQ) [39] and GKAPR5 (PDB ID: 6TQ0) [39]. A similar conformation is present in the complex between E5 and Arc-NL.

### Simultaneous binding of anti-Arc-NL and anti-Arc-CL nanobodies to full-length Arc

Our earlier data, including crystal structures, showed that the nanobodies H11 and C11 independently bound to the Arc-NL and Arc-CL, respectively, at the same time [37]. Therefore, as H11 and E5 both bind to the same pocket in Arc-NL, we wished to confirm if this is also true for the nanobody pair E5 and C11.

As a control, we carried out analytical SEC of the Arc-CTdt construct with H11 and C11, i.e. the complex crystallized previously. As expected, the SEC peak for the Arc dimer moves when C11 and H11 are sequentially added (Fig 7A).

**Fig 7.**
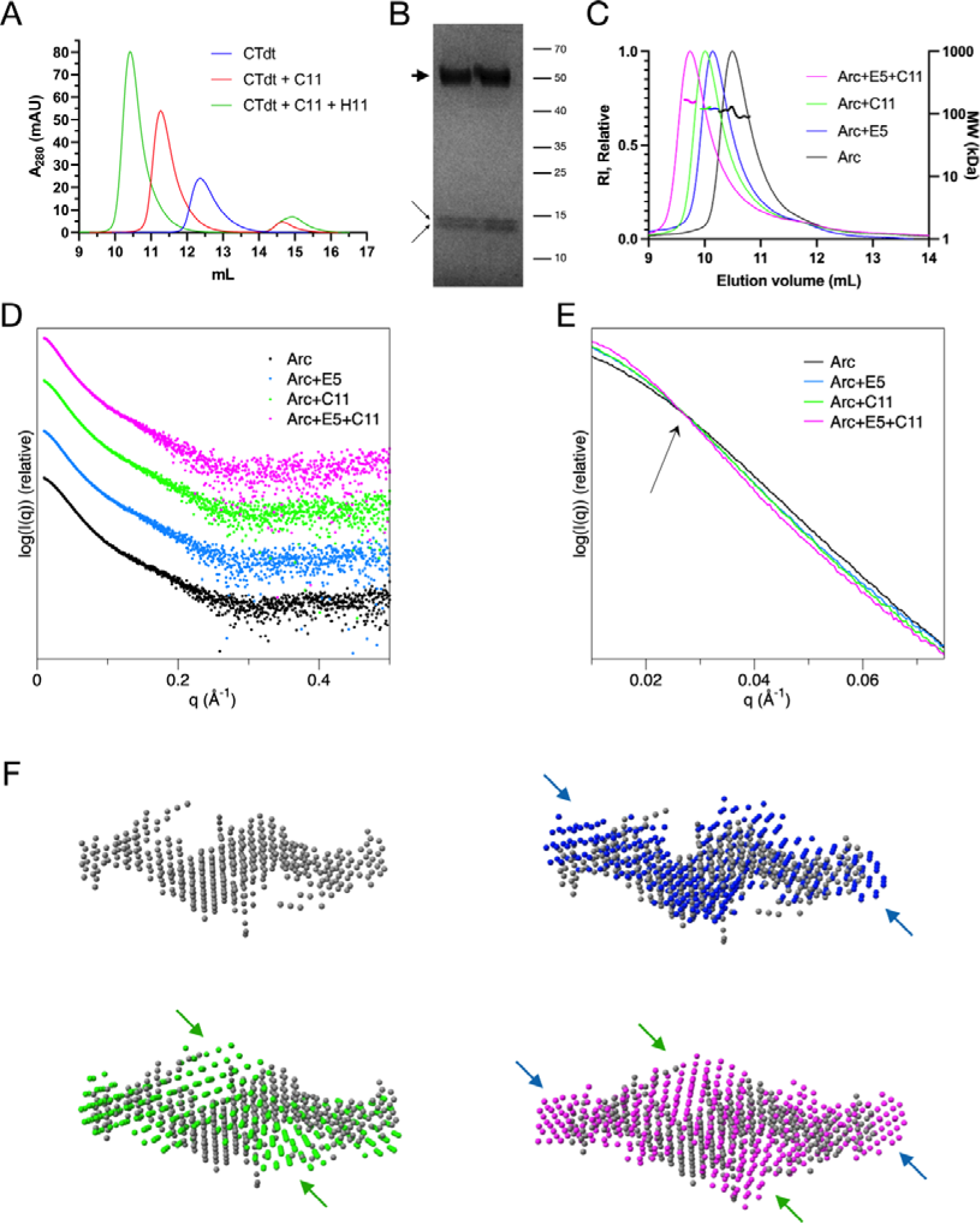
Binding of two nanobodies simultaneously. A. Analytical SEC of CTdt with nanobodies C11 and H11 indicates simultaneous binding. B. SDS-PAGE of purified ternary complex rArcFL-7A+E5+C11. Arc is shown with the thick arrow and the two nanobodies with thin arrows. C. SEC-MALS of rArcFL-7A with nanobodies E5 and C11. D. SAXS data; displaced along the y axis for clarity. E. Zoom-in of the low-angle region indicates different shapes for the scattering curves, with an increase in size upon addition of each nanobody. The crossover point is indicated by an arrow. The SAXS curves were scaled together for the analysis. F. Ab initio dummy atom models for Arc (gray), Arc+E5 (blue), Arc+C11 (green), Arc+E5+C11 (magenta). Apparent positions of extra density upon nanobody addition are shown with arrows.

We purified the flArc dimeric mutant rArcFL-7A in complex with both E5 and C11, and we used the samples for low-resolution structural studies in solution using SEC-SAXS, as well as for molecular weight determination in SEC-MALS. In the purified complex, bands for both nanobodies can be detected in addition to flArc (Fig 7B), indicating ternary complex formation between the three proteins. In the SAXS and MALS experiments (Table 6, Fig7C-F), the dimeric state of flArc was confirmed, and the addition of either E5 or C11 caused an expected increase in MW. This increase was effectively doubled, when both nanobodies were added. This shows simultaneous binding of both E5 and C11 to flArc, similarly to the pair H11-C11, for which we have high-resolution data from before with the Arc-CTdt [37].

**Table 6.** SAXS and SEC-MALS parameters. All data point towards the binding of E5 and H11 to the Arc dimer simultaneously, resulting in a 2:2:2 ternary complex.

| **Complex** | **rArcFL7A** | **+E5** | **+C11** | **+E5 and C11** |
| --- | --- | --- | --- | --- |
| *Oligomeric state/*<br>*stoichiometry* | Dimer | 2:2 | 2:2 | 2:2:2 |
| *Theoretical mass (kDa)* | 90.0 | 115.6 | 116.2 | 141.7 |
| *Mass from SEC-MALS*<br>*(kDa)* | 104.9 | 122.0 | 123.4 | 163.0 |
| *Mass from SAXS*<br>*Bayesian estimate (kDa)* | 109.1 | 130.1 | 138.2 | 157.1 |
| *Guinier $R_g$ (Å)* | 47.4 | 55.8 | 57.2 | 59.8 |
| *$P(r)$ $R_g$ (Å)* | 50.5 | 58.5 | 56.6 | 63.7 |
| *$D_{max}$ (Å)* | 200 | 228 | 211 | 236 |
| *Porod volume (nm<sup>3</sup>)* | 230 | 298 | 294 | 350 |

SAXS was used to assess the binding sites of the nanobodies and thus, the likely location of the Arc lobe domains in the flArc dimer. The scattering curves (Fig 7D-E, Table 6) indicate that rArcFL7A is a dimer and that E5 and H11 can bind simultaneously in a 2:2:2 stoichiometry. Considering the current models (Fig 7F) and earlier data from complexes of Arc with nanobodies E5 and H11 (binding to the NL) and C11 (binding to the CL) [37, 38], one can conclude that in the dimeric full-length Arc, the N-lobe is close to the ends of the elongated dimer, and the C-lobe is more centrally located. The dimer interface is formed by the NTD [37].

### Insights into potential binding partners of Arc-NL

To search for potential binding partners of Arc-NL, the CDR3 sequences of both E5 and H11 were used for database searches. The search was restricted to human, mouse, and rat protein sequences. For each CDR3, an interesting match was noted (see below). Several other matches were considered, and some of them are collected in S1 Table. Such a short sequence gives many hits, especially if one looks at the Arc-NL binding motif alone.

For the E5 CDR3, all selected hits show the consensus motif PxWS2(Fig); note that E5 is the only currently known ligand of Arc-NL with a Trp residue in the aromatic position. Therefore, it would be interesting to find possible endogenous ligands with the same motif. The closest hit was cathepsin H. Cathepsins are proteases targeted to endosomes and lysosomes, associated with the active regulation of synaptic plasticity [71]. Cathepsin H can regulate neuropeptide-associated signaling cascades; it is mainly expressed in astrocytes and found at high concentrations in cerebral spinal fluid [70]. As capsid-transferred neuronal Arc mRNA has been observed in astrocytes [73, 74], activity-dependent interaction between cathepsin H and Arc could be tested as a hypothesis.

Regarding the short-listed hits for the H11 CDR3, only the PRKCA-binding protein displays the PxY consensus motif (S2 Fig); the sequence corresponds to the protein PICK1 [75]. PICK1 is involved in the endocytosis of AMPAR and interacts directly with Arc through its BAR domain (amino acids 152-362) [76, 77]. Fig 8 shows homology between the H11 CDR3 and PICK1 amino acids 278-294 in the BAR domain; this segment corresponds to a flexible loop at the end of the BAR domain and would therefore be well accessible for Arc-NL binding. Thus, the H11 nanobody CDR3 loop may have helped in identifying the specific Arc interaction site on PICK1. This hypothesis can be tested in further experiments to reveal molecular details of the Arc-PICK1 interaction.

**Fig 8.**
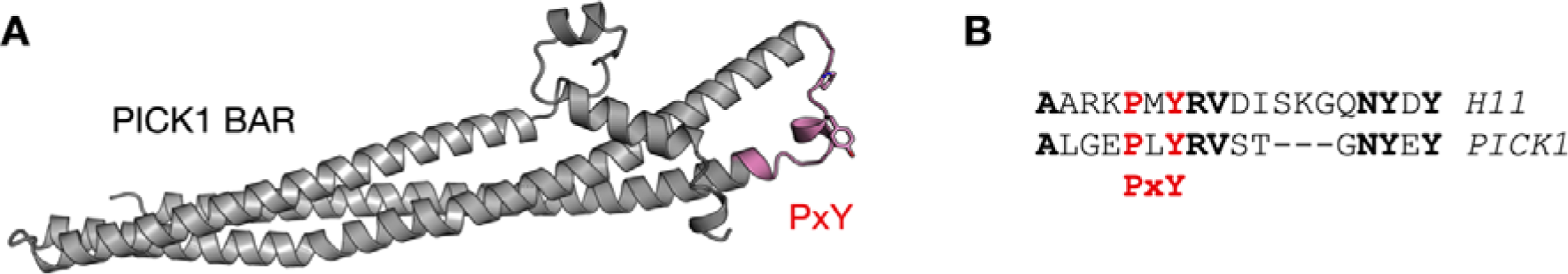
Identification of a putative Arc-NL binding site on PICK1. A. AlphaFold2 model of the PICK1 BAR domain shows that the distal loop carries a PxY motif (pink). The side chains of the Pro and Tyr residues in the motif are shown. B. Sequence alignment between the H11 CDR3 loop and the PxY motif of PICK1. Bold residues are identical, and the PxY motif is highlighted in red.

### Conclusions

Nanobodies are emerging as versatile molecular tools for both basic and translational research. We characterized the structural and functional properties of two nanobodies (H11 and E5) against Arc, a complex molecular regulator of synaptic plasticity. Our results pave the way towards the use of anti-Arc nanobodies H11 and E5 as molecular modulators of Arc *in vitro* and *in vivo,* possibly in combination with nanobodies simultaneously targeting the Arc-CL. Structural biology of different oligomeric forms of Arc will benefit from the use of different nanobody combinations. Besides, nanobodies can be delivered to living cells *in vitro* and have been used in super-resolution microscopy, highlighting nanobodies as diagnostic and functional tools [78, 79]. Furthermore, several strategies are being developed to promote the use of nanobodies in the treatment and diagnosis of brain diseases *in vivo* [80–82]. For example, nanobodies have been successfully used to treat an animal model of Alzheimer’s disease [83]. Considering the latest advances in nanobody-associated research, the use of anti-Arc nanobodies for behavioral, diagnostic, and therapeutic purposes is within range of current translational neuroscience. Anti-Arc nanobodies E5 and H11 provide molecular tools to selectively study how Arc structure, function, and oligomerization influence memory, learning, stress regulation, and neuropathologies.

## Acknowledgements

We are grateful to Ju Xu and Anne Baumann for technical assistance during this study, and we acknowledge the use of the Core Facility for Biophysics, Structural Biology, and Screening (BiSS) at the University of Bergen, which has received infrastructure funding from the Research Council of Norway through NORCRYST (grant number 245828) and NOR– OPENSCREEN (grant number 245922). We thank DESY (Hamburg, Germany) for the provision of their experimental facilities; part of this research was carried out on the PETRA III storage ring. Our gratitude extends to the ISA facility (Aarhus, Denmark), as part of this research was conducted on the storage ring ASTRID2. We further recognize access to and support at the SOLEIL and MAX-IV synchrotrons.

## Supporting information

**S1 Fig.** ITC assays for Arc-nanobody interactions (individual replicates).

**S2 Fig.** Sequence alignments between CDR3 sequences and the top hits.

**S1 Table.** Top hits from the nanobody CDR3 BLAST search.

## Notes

### Competing Interest Statement

The authors have declared no competing interest.

