## Supplementary material for "Structural characterization of two nanobodies targeting the ligand-binding pocket of human Arc": S1 Table

***S1 Table. Protein sequence matches for the CDR3s of E5 and H11 provided by Blast****. For each sequence, its corresponding protein, species, E-value and ID are shown. Relevant matches (E < 10) are highlighted in bold. Abbreviations: HCCA2, hepatocellular carcinoma-associated protein 2; PRKCA, protein kinase C alpha; TFIID, transcription factor II D.*

| **Query** | **Protein match** | **Species** | **E-value** | **Sequence ID** |
|---|---|---|---|---|
| **E5-CDR3** | Pro-cathepsin H | *Homo sapiens* | 3.4 | 6CZK (PDB) |
| | Cathepsin H | *Homo sapiens* | 3.4 | AAH02479.1 |
| | HCCA2 | *Homo sapiens* | 28 | BAB62266.1 |
| | TFIID – Subunit 2 | *Rattus norvegicus* | 81 | NP_579853.1 |
| | TFIID – Subunit 2 (Isoforms X1 and X2) | *Mus musculus* | 81 | XP_011243959.1 |
| **H11-CDR3** | Perinuclear binding protein | *Mus musculus* | 0.98 | CAA86675.1 |
| | Poly(U)-specific endoribonuclease precursor | *Rattus norvegicus* | 16 | NP_001177998.1 |
| | Unconventional myosin XV | *Mus musculus* | 16 | 7UDT (PDB) |
| | PRKCA-binding protein | *Homo sapiens* | 31 | NP_001034672.1 |
| | PRKCA-binding protein | *Rattus norvegicus* | 31 | NP_445912.2 |
| | PRKCA-binding protein | *Mus musculus* | 31 | NP_001039023.1 |
