## Supplementary material for "Structural characterization of two nanobodies targeting the ligand-binding pocket of human Arc": S2 Fig

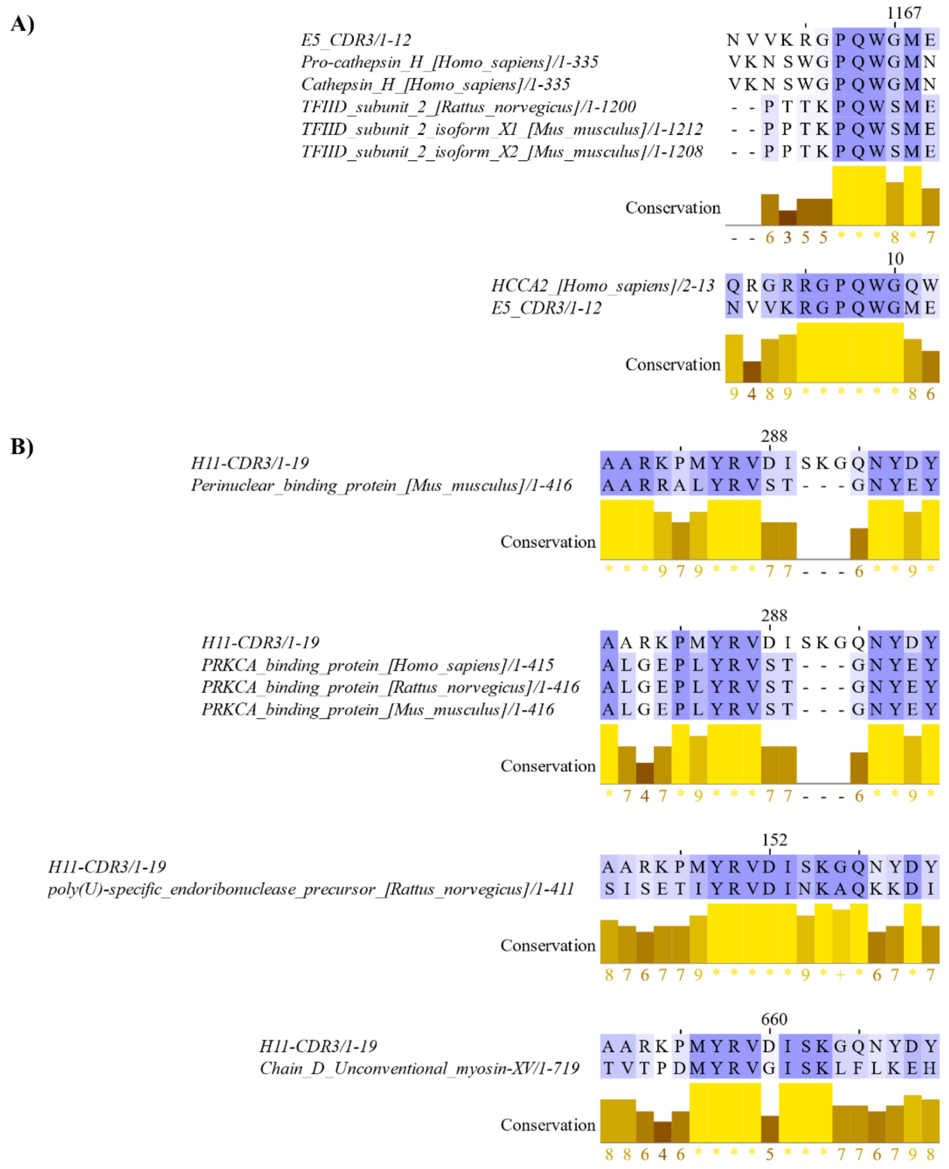


**S2 Fig. Multiple sequence alignment using Blast hits for E5 and H11 CDR3 loops.** (A) Multiple sequence alignment using Clustal Omega [61] of the E5 CDR3 with the protein sequences obtained in Blast [60]. (B) Pairwise alignment performed with Clustal Omega of the H11 CDR3 with the protein sequences acquired from Blast. The data were visualized with Jalview.
