## Supplementary figures and images for "Structural characterization of two nanobodies targeting the ligand-binding pocket of human Arc"

### S1 Fig

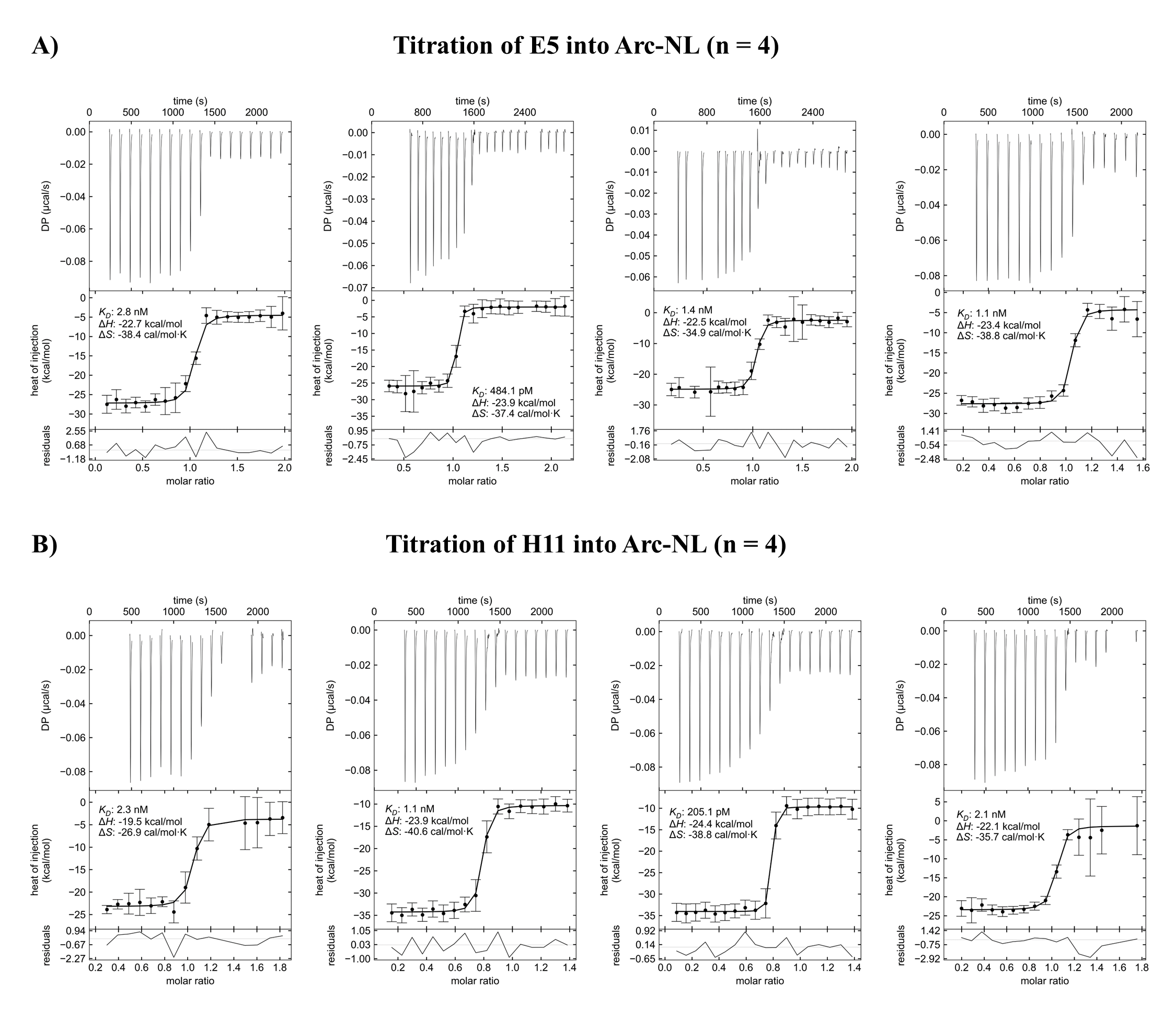
